# Metabolomic profiling identifies complex lipid species associated with response to weight loss interventions

**DOI:** 10.1101/2020.10.05.326025

**Authors:** Nathan A. Bihlmeyer, Lydia Coulter Kwee, Clary B. Clish, Amy Anderson Deik, Robert E. Gerszten, Neha J. Pagidipati, Blandine Laferrère, Laura P. Svetkey, Christopher B. Newgard, William E. Kraus, Svati H. Shah

**Affiliations:** Duke Molecular Physiology Institute, Duke University, Durham, NC 27701, USA; Metabolomics Platform, Broad Institute of MIT and Harvard, Cambridge, MA 02142, USA; Division of Cardiovascular Medicine, Beth Israel Deaconess Medical Center, Harvard Medical School, Boston, MA 02215, USA; Duke Clinical Research Institute, Duke University, Durham, NC 27701, USA; Columbia University Irving Medical Center, New York, NY 10032, USA; Department of Medicine, Duke University Medical Center, Durham, NC 27710, USA

## Abstract

Obesity is an epidemic internationally. While weight loss interventions are efficacious, they are compounded by heterogeneity with regards to clinically relevant metabolic responses. Thus, we sought to identify metabolic pathways and biomarkers that distinguish individuals with obesity who would most benefit from a given type of intervention. Liquid chromatography mass spectrometry-based profiling was used to measure 765 metabolites in baseline plasma from three different weight loss studies: WLM (behavioral intervention, N=443), STRRIDE-PD (exercise trial, N=163), and CBD (surgical cohort, N=125). The primary outcome was percent change in insulin resistance (as measured by the Homeostatic Model Assessment of Insulin Resistance [%ΔHOMA-IR]) over the intervention. Overall, 92 individual metabolites were associated with %ΔHOMA-IR after adjustment for multiple comparisons. Concordantly, the most significant metabolites were triacylglycerols (TAGs; p=2.3e-5) and diacylglycerols (DAGs; p=1.6e-4), with higher TAG and DAG levels associated with a greater improvement in HOMA-IR. In tests of heterogeneity, 50 metabolites changed differently between weight loss interventions; we found amino acids, peptides, and their analogues to be most significant (4.7e-3) in this category. Our results highlight novel metabolic pathways associated with heterogeneity in response to weight loss interventions, and related biomarkers which could be used in future studies of personalized approaches to weight loss interventions.

## Introduction

Obesity is a major epidemic in the developed world and is an increasing problem in developing countries, with a range of consequences including dyslipidemia, hypertension, cardiovascular disease (CVD), stroke, type 2 diabetes mellitus (T2DM), and overall mortality.[1–4] In the United States, one third of adults are obese and approximately 300,000 deaths are attributed to obesity every year.[5,6] Behavioral, pharmacologic, exercise, dietary, and surgical intervention methods have been attempted to curb the obesity epidemic. Ideally, these interventions would be effective in all adults equally; however even when accounting for the amount of weight loss, individuals show heterogeneity in improvement in obesity-related co-morbidities and CVD risk factors.[7] Further, compliance with obesity intervention protocols is low unless expensive and protracted intervention programs run by a trained interventionist are employed. Surgical obesity interventions can partially overcome compliance issues; however, these are costly, can have short- and long-term complications, and are characterized by frequent weight regain. In recognition of these issues, the American Heart Association (AHA), the American College of Cardiology (ACC), and The Obesity Society (TOS) released a call for researchers to focus on determining the “the best approach to identify and engage those who can benefit from weight loss”.[8]

Blood-based biomarkers, by serving as more granular snapshots into an individual’s unique biochemistry, could distinguish individuals who would benefit the most from surgical interventions from those who can benefit from lower cost, less-invasive solutions. Our previous work has demonstrated a clear disconnect between amount of weight loss during lifestyle obesity interventions and improvement in insulin resistance[9], as well as marked inter-individual heterogeneity in amount of weight loss and metabolic responses to a given weight loss intervention. In this study, we investigate an omics-driven personalized approaches to understanding the molecular mechanisms of obesity and identifying biomarkers of response among diverse weight loss interventions including behavioral, exercise and surgical interventions.

## Methods

### Study Populations and Study approval

The **WLM** cohort has been described previously[10] (NCT00054925); all participants provided written consent. Approval from the Duke University Institutional Review Board was given. Notably, overweight or obese adults with hypertension, dyslipidemia, or both were recruited from clinical research centers at Duke University, Johns Hopkins University, Pennington Biomedical Research Center, or the Kaiser Permanente Center for Health Research between 2003 and 2009. This study only involves samples collected during phase 1 of the WLM intervention in which all participants were involved in a group-based behavioral intervention. A trained interventionist led 20 weekly group sessions with the goals for participants to reach 180 minutes per week of moderate physical activity (e.g., walking), reduce caloric intake, adopt the Dietary Approaches to Stop Hypertension (DASH) dietary pattern, and lose approximately ~0.5-1 kg per week. DASH was chosen because it reduces CVD risk factors.[11–16] Only participants who lost at least 4 kg during the 6-month weight loss program were considered for this study. Relevant for this study, blood plasma was drawn at baseline and 6-month follow-up, which we used for non-targeted metabolomics.

The **STRRIDE-PD** study[17,18] (NCT00962962) compared three 6-month exercise-only groups between 2009 and 2013; differing in amount or intensity to a fourth lifestyle intervention group: diet plus exercise similar to the first 6-months of the Diabetes Prevention Program (DPP). All participants in STRRIDE-PD provided written consent and approval from the Duke University Institutional Review Board was given. Sedentary, moderately overweight/obese (25 < BMI < 35 kg/m^2^), nonsmoking adults between the ages of 45 and 75 years with prediabetes, but no history of diabetes mellitus (T2DM) or CVD, were randomly assigned to one of four groups. The STRRIDE-PD study defined prediabetes as high-normal to impaired fasting glucose (95-125 mg/dL). The four groups were: 1) low-amount/moderate-intensity exercise; 2) high-amount/moderate-intensity exercise; 3) high-amount/vigorous-intensity exercise; 4) diet and exercise clinical lifestyle intervention. Relevant for this study, the blood plasma which we used for non-targeted metabolomics was drawn at baseline and 6-month follow-up.

The **CBD** cohort was a surgical weight loss cohort of individuals who underwent either Roux-en-Y gastric bypass surgery (RYGB) or adjustable gastric banding surgery (AGB) at St Luke’s Roosevelt Hospital Center between 2006 and 2014. All participants signed a consent form prior to engaging in various research studies aiming to identify changes in gut hormones and metabolism after bariatric surgery (NCT01516320, NCT02287285, NCT02929212, NCT00571220) [19–23]. Approval from the Columbia University Institutional Review Board was given. As such, each participant had fasting blood plasma drawn at baseline, with body weight and HOMA-IR measured at multiple follow-up visits up to a year.

#### Metabolomic profiling

Four complementary liquid chromatography tandem mass spectrometry (LC-MS)-based methods were used to measure plasma lipids and polar metabolites as previously described [24–28]. The methods are characterized by the chromatography stationary phase and MS ionization mode used and are referred to as C8-pos, C18-neg, HILIC-pos, and HILIC-neg. The C8-pos, C18-neg, and HILIC-pos methods were configured on LC-MS systems comprised of Nexera X2 U-HPLCs (Shimadzu) coupled to Q Exactive series orbitrap mass spectrometers (Thermo Fisher Scientific) for high resolution accurate mass (HRAM) profiling of both hundreds of identified metabolites and thousands of unknowns, while the HILIC-neg method was operated on both a Nexera X2-Q Exactive system for HRAM profiling and a UPLC (Waters) coupled to a QTRAP 5500 (SCIEX) for targeted profiling.

The **C8-pos** method measures polar and nonpolar lipids. Lipids were extracted from 10 μL plasma using 190 μL of isopropanol, separated using reversed phase C8 chromatography, and analyzed HRAM, full-scan MS in the positive ion mode. The **C18-neg** method measures free fatty acids, oxidized fatty acids and lipid mediators, bile acids, and metabolites of intermediate polarity; these metabolites were extracted from 30 μL plasma using 90 μL of methanol, then separated using reversed phase C18 chromatography, and analyzed using HRAM, full-scan MS in the negative ion mode. The **HILIC-pos** method measures amino acids, amino acid metabolites, acylcarnitines, dipeptides, and other cationic polar metabolites; these metabolites were extracted from 10 μL plasma using 90 μL of 25% methanol/75% acetonitrile, then separated using hydrophilic interaction liquid chromatography (HILIC), and analyzed using HRAM, full-scan MS in the positive ion mode. The **HILIC-neg** method measures sugars, organic acids, purines, pyrimidines, and other anionic polar metabolites; these metabolites were extracted from 30 μL plasma using 120 μL of methanol containing internal standards and analyzed using either HRAM, full-scan MS in the negative ion mode or targeted multiple reaction monitoring using the QTRAP 5500 triple quadruple MS system.

Raw data from Q Exactive series mass spectrometers were processed using TraceFinder software (Thermo Fisher Scientific) to detect and integrate as subset of identified metabolites and Progenesis QI software (v 2.0, Nonlinear Dynamics) to detect, de-isotope, and integrate peak areas from both identified and unknown metabolites. MultiQuant (SCIEX) was used to integrate peak areas of metabolites measured using the QTRAP 5500. Identities of metabolites were confirmed by matching measured retention times (RT) and mass-to-charge ratios (m/z) to authentice reference standards. Since reference standards are not available for all lipids, representative lipids from each lipid class were used to characterize RT and m/z ratio patterns. Lipid identities are reported at the level of lipid class, total acyl carbon content, and total double bond content since the LC-MS method does not discretely resolve all isomeric lipids from one another. Unknown features (unnamed metabolites) were not used in these analyses. Coefficients of variation (CVs) and missingness are reported in Supplemental Tables 1 & 2 for each metabolite.

#### Statistical analysis

Percent change in Homeostatic Model Assessment of Insulin Resistance (HOMA-IR) over the intervention time period is the outcome used to represent metabolic health for this study. HOMA-IR is calculated from clinically determined blood glucose and insulin levels as previously described.[10,11] Individuals were excluded if HOMA-IR percent change was implausible, i.e., greater than five standard deviations from the cohort mean (N=2 removed). Metabolites measured at baseline in blood plasma by LC-MS are in the form of the natural log of the MS peak’s area under the curve (AUC). The primary study was association of baseline metabolites with percent change in HOMA-IR over the obesity intervention time period. The secondary study was of heterogeneity between cohorts in the primary study.

### Within cohort analysis

Two statistical analyses were performed for this study within each of the three cohorts. 1) A univariate linear association model between percent change of HOMA-IR and baseline metabolite, in order to find metabolites that effect HOMA-IR. 2) The same model as before with the addition of covariates for age, sex, race, baseline clinical triglycerides, and percent change in weight over the follow-up time period in order to determine if the individual metabolite effects HOMA-IR independent of traditional clinical measures known to be associated with insulin resistance.

### Meta-analysis

For both the primary univariate model and the full model, the three cohorts then were meta-analyzed together using an inverse-variance weighted random effects model implemented in R’s meta library.[29] To account for the multiple tests, a false discovery rate (FDR) correction was applied to the random effects p-values from the meta-analysis of the three cohorts.[30] A Cochran q-test of heterogeneity also was performed to find metabolites having different effects between the three cohorts.[31] Metabolites were filtered for only known metabolites with less than 25% missingness after meta-analysis (N=199 removed).

### Metabolite set analysis

We used a variation of the Gene Set Enrichment Analysis (GSEA) method [32,33] to determine in an unbiased manner if any particular group of metabolites was overrepresented at the beginning or end of a sorted list of our 765 metabolite results. Lists of metabolite results were either sorted by z-score from the random effects meta-analysis (primary analysis) or by Cochran q-test statistic from the heterogeneity analysis (secondary analysis). The goal of GSEA is to determine whether the metabolites are randomly distributed throughout this sorted list or mainly found at the beginning or end by performing a random walk along the list and recording the maximum enrichment score (ES) along the way. Metabolite sets associated with the outcome will have positive or negative ES depending on the direction of effect; while metabolite sets unrelated to the outcome will have ES near zero. The 22 sets from HMDB’s taxonomic sub-classification of metabolites were used, with only sets with at least five metabolites in our data were tested. P-values were determined by one million permutations and FDR multi-test correction.

## Results

### Study Populations

This study includes three intervention study cohorts: Weight Loss Maintenance (WLM) cohort, Studies of Targeted Risk Reduction Interventions through Defined Exercise in individuals with Pre-Diabetes (STRRIDE-PD), and Columbia Bariatric and Diabetes (CBD) cohort. Table 1 describes the demographics of these three cohorts.

**Table 1:** Baseline Characteristics of the Study Populations.

| Clinical Characteristics | WLM | STRRIDE-PD | CBD |
| --- | --- | --- | --- |
| N | 443 | 163 | 125 |
| Female (%) | 62% | 63% | 82% |
| Race (% AA) | 37% | 18% | 50% |
| Race (% EA) | 62% | 78% | 49% |
| Age (years) | 56 ± 8.7 | 59 ± 7.5 | 42 ± 10 |
| Weight (kg) | 96 ± 16 | 86 ± 12 | 121 ± 22 |
| Weight Loss (kg) | -8.5 ± 3.8 | -2.7 ± 4.0 | -35 ± 16 |
| BMI (kg/m<sup>2</sup>) | 34 ± 4.7 | 30 ± 2.8 | 44 ± 5.8 |
| HOMA-IR | 2.4 ± 1.6 | 2.0 ± 1.5 | 9.6 ± 5.8 |
| HOMA-IR, Percent Change (%) | -16 ± 105 | -16 ± 42 | -71 ± 21 |
| Triglycerides (mg/dL) | 127 ± 71 | 123 ± 64 | 158 ± 108 |
| Total Cholesterol (mg/dL) | 185 ± 35 | 190 ± 32 | 187 ± 39 |
| HDL Cholesterol (mg/dL) | 43 ± 14 | 46 ± 14 | 48 ± 12 |
| LDL Cholesterol (mg/dL) | 110 ± 27 | 119 ± 27 | 108 ± 31 |

Unless otherwise stated, measures are at baseline timepoint. AA: African American. EA: European Ancestry. BMI: body mass index. HOMA-IR: Homeostatic Model Assessment of Insulin Resistance. HDL: high density lipoprotein cholesterol. LDL: low density lipoprotein cholesterol. Weight loss is measured over the follow-up period. Values are listed as mean ± standard deviation. “cm” is centimeters. “kg” is kilograms. “m” is meters. “mg” is milligrams. “dL” is deciliters. “%” is percent.

### Metabolomic profiling

Four complementary liquid chromatography tandem mass spectrometry (LC-MS)-based methods were used to measure plasma lipids and polar metabolites. Figure 1 shows the Human Metabolome Database’s (HMDB’s)[34–37] superclass and subclass taxonomic identifications for the 765 known metabolites that were identified from all LC-MS methods in this study. Note that HMDB IDs ascribed to lipids are representative of one or more isomers sharing the same chemical formula.

**Fig. 1:**
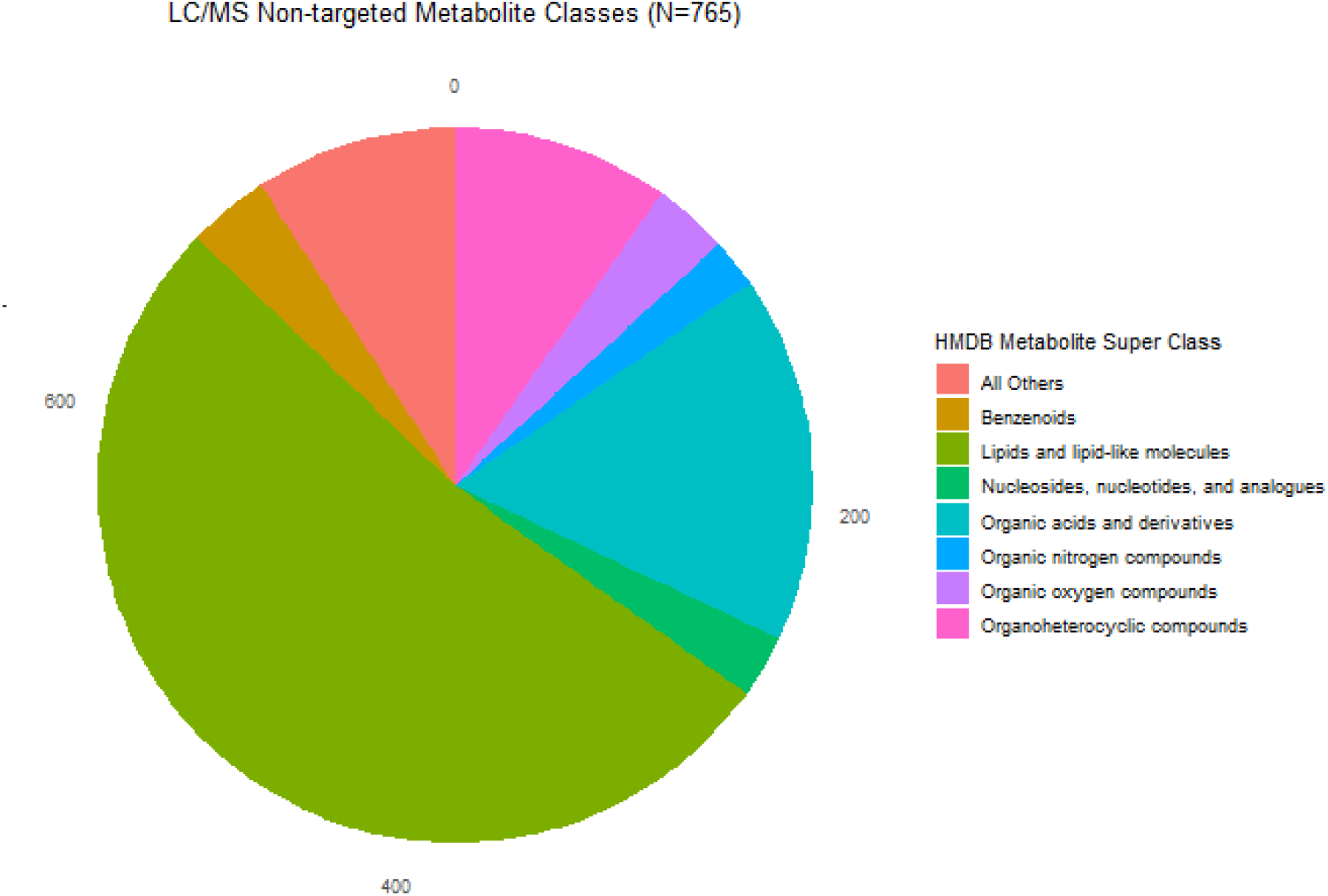
LC-MS Non-targeted Metabolite Classes.

Pie graph of the Human Metabolome Database’s (HMDB’s) superclass taxonomic identifications for the 765 known metabolites from all LC-MS methods in this study. Nearly half of known metabolites were lipids or lipid-like molecules shown in forest green color.

### Association of baseline metabolites with percent change in HOMA-IR

Meta-analysis of the univariate linear regression models in the three weight loss intervention cohorts identified 92 baseline metabolites (Figure 2, Supplemental Table 1) associated with percent change in HOMA-IR after adjustment for multiple comparisons (FDR<5%). The top metabolites were dominated by triacylglycerol (TAG) and diadylglycerol (DAG) species (in light-orange and red, respectively, in Figure 2) and are listed in Table 2. As evidenced by the negative beta coefficient, higher baseline levels of metabolites at baseline were associated with greater reduction in HOMA-IR. For instance, the top result is C34:0 DAG with a FDR adjusted p-value of 5.42e-6 and an effect size of −36.08% change in HOMA-IR over the intervention. To determine if an individual metabolite had an effect on HOMA-IR independent of traditional clinical measures known to be associated with insulin resistance, we looked for nominal significance in the full model. In the full model adjusted for age, sex, race, baseline triglycerides, and percent change in weight over the follow-up time period, 90 of the 92 significant metabolites retained the same direction of effect, 38 of these remained nominally associated with percent change in HOMA-IR (Supplemental Table 2). Table 2 compares the effect size estimates of the univariate model with the full model for the top 10 metabolites, demonstrating significance and consistency of magnitude and directionality of effect.

**Fig. 2:**
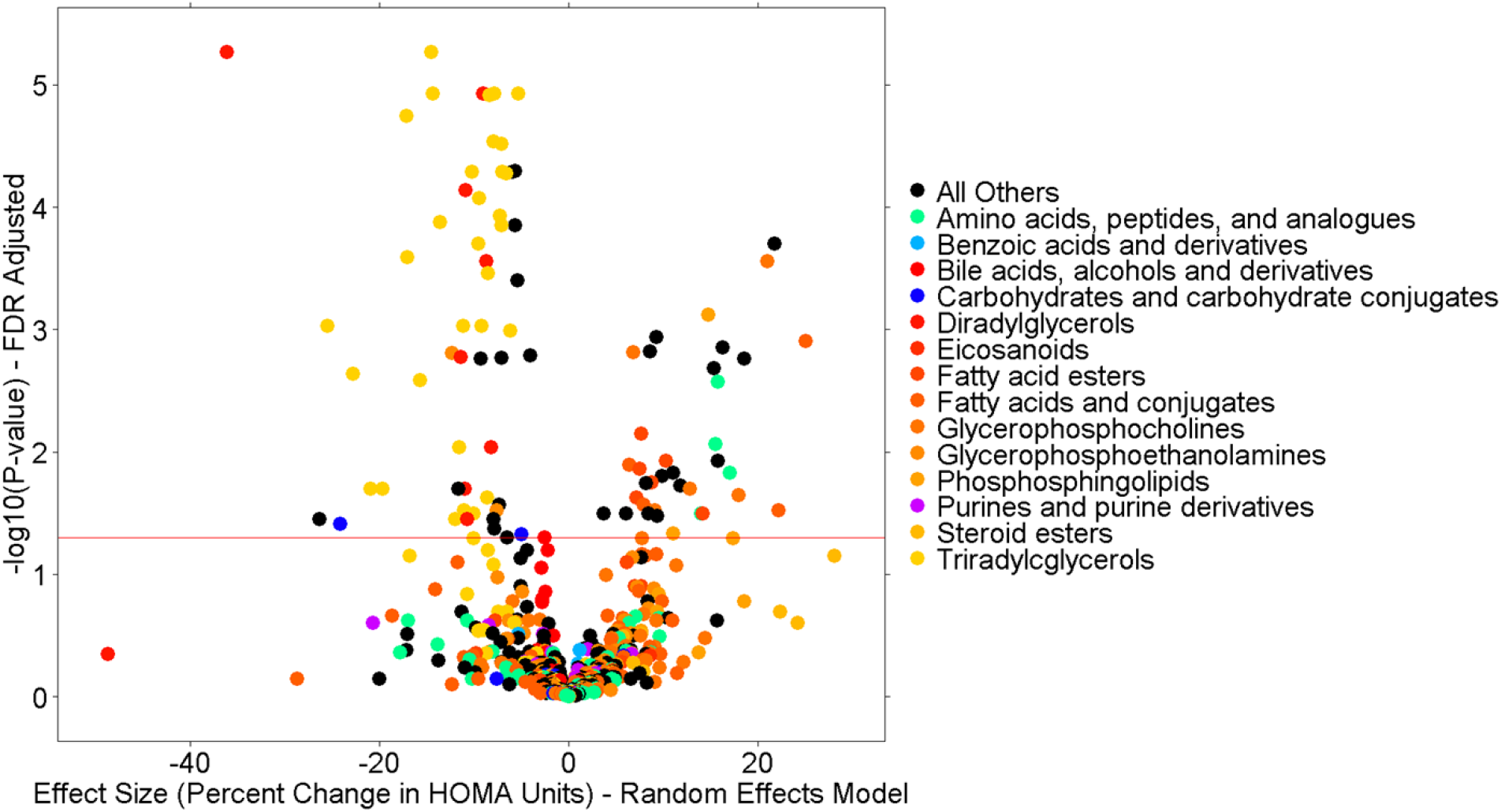
Meta-analysis of the Univariate Model.

**Table 2:** Comparison of effect sizes between univariate and full models.

| Metabolite | Univariate Model |  |  |  | Full Model |  |  |
| --- | --- | --- | --- | --- | --- | --- | --- |
|  | Pvalue_FDR | N | Pvalue | Beta | Beta | Pvalue | N |
| C34:0 DAG | 5.42E-06 | 580 | 1.44E-08 | -36.08 | -25.03 | 8.90E-03 | 571 |
| C50:1 TAG | 5.42E-06 | 677 | 1.92E-08 | -14.55 | -9.21 | 4.42E-03 | 661 |
| C30:0 DAG | 1.19E-05 | 676 | 8.83E-08 | -9.02 | -5.26 | 1.67E-01 | 660 |
| C43:1 TAG | 1.19E-05 | 662 | 9.07E-08 | -5.35 | -3.37 | 1.80E-01 | 646 |
| C48:0 TAG | 1.19E-05 | 677 | 1.26E-07 | -7.80 | -4.79 | 2.32E-03 | 661 |
| C50:2 TAG | 1.19E-05 | 677 | 1.16E-07 | -14.34 | -8.11 | 2.87E-02 | 661 |
| C56:1 TAG | 1.23E-05 | 677 | 1.52E-07 | -8.31 | -6.23 | 5.36E-02 | 661 |
| C51:2 TAG | 1.81E-05 | 677 | 2.56E-07 | -17.08 | -11.22 | 1.71E-03 | 661 |
| C45:2 TAG | 2.91E-05 | 677 | 4.63E-07 | -7.89 | -4.53 | 2.51E-02 | 661 |
| C46:1 TAG | 3.06E-05 | 677 | 5.40E-07 | -7.09 | -3.69 | 3.82E-02 | 661 |

Comparison of univariate model individual metabolite analysis (the primary analysis) to the full model conditional analyses where traditional clinical measures of age, sex, race, baseline clinical triglycerides, and percent change in weight over the follow-up time period were added to the model. The lack of major changes in effect sizes between the two models indicates the individual metabolites identified in the primary analysis provide additional clinical value over the traditional clinical measures. “Metabolite” is the common name for the metabolite. “N” is the number of individuals in the meta-analysis model. “Pvalue” is the random-effects p-value for the model. “Pvalue_FDR” is the primary p-value after FDR multi-test correction. “Beta” is the random-effects effect size estimate for the model. Pvalues are colored green if < 0.05 and Betas and colored blue to red for being high or low, respectively.

Volcano plot of the main analysis results from meta-analysis of the three cohorts for the 765 known metabolites from all LC-MS methods in this study. The x-axis is the random-effects effect-size estimate in percent change of HOMA-IR units (e.g., “−40” indicates a 40% drop in HOMA-IR over the intervention time period). Each dot is a metabolite colored by the Human Metabolome Database’s (HMDB’s) subclass taxonomic identification. The y-axis is the – log10(P-value) from the random effects meta-analysis of the three cohorts in this study after FDR correction.

### Metabolite set analysis to identify enriched biological pathways

We used metabolite set analysis to determine in an unbiased manner if any particular group of metabolites was overrepresented within the HMDB’s taxonomic sub-classification of metabolites. Figure 3 shows the enrichment plots for the top five sub-classification of metabolites most associated with increased in insulin resistance followed by the top five sub-classification of metabolites most associated with reduction in insulin resistance. TAGs (FDR adjusted p=2.3e-5) and DAGs (FDR adjusted p=1.6e-4) were the most significant sub-classifications associated with reduction in insulin resistance over the intervention period in all 3 cohorts after FDR correction for the 22 sub-classifications tested. In fact, both TAGs and DAGs are associated with reduction in insulin resistance over the intervention period in this regard. The “Glycerophosphocholines” (FDR adjusted p=3.8e-4), “Fatty acid esters” (FDR adjusted p=1.3e-3), and “Phosphosphingolipids” (FDR adjusted p=1.1e-2) sub-classifications were also significantly associated with insulin resistance after FDR multi-test correction; however, they were associated with increased insulin resistance over the intervention period. Supplemental Table 3 contains results from all tested HMDB sub-classifications.

**Fig. 3:**
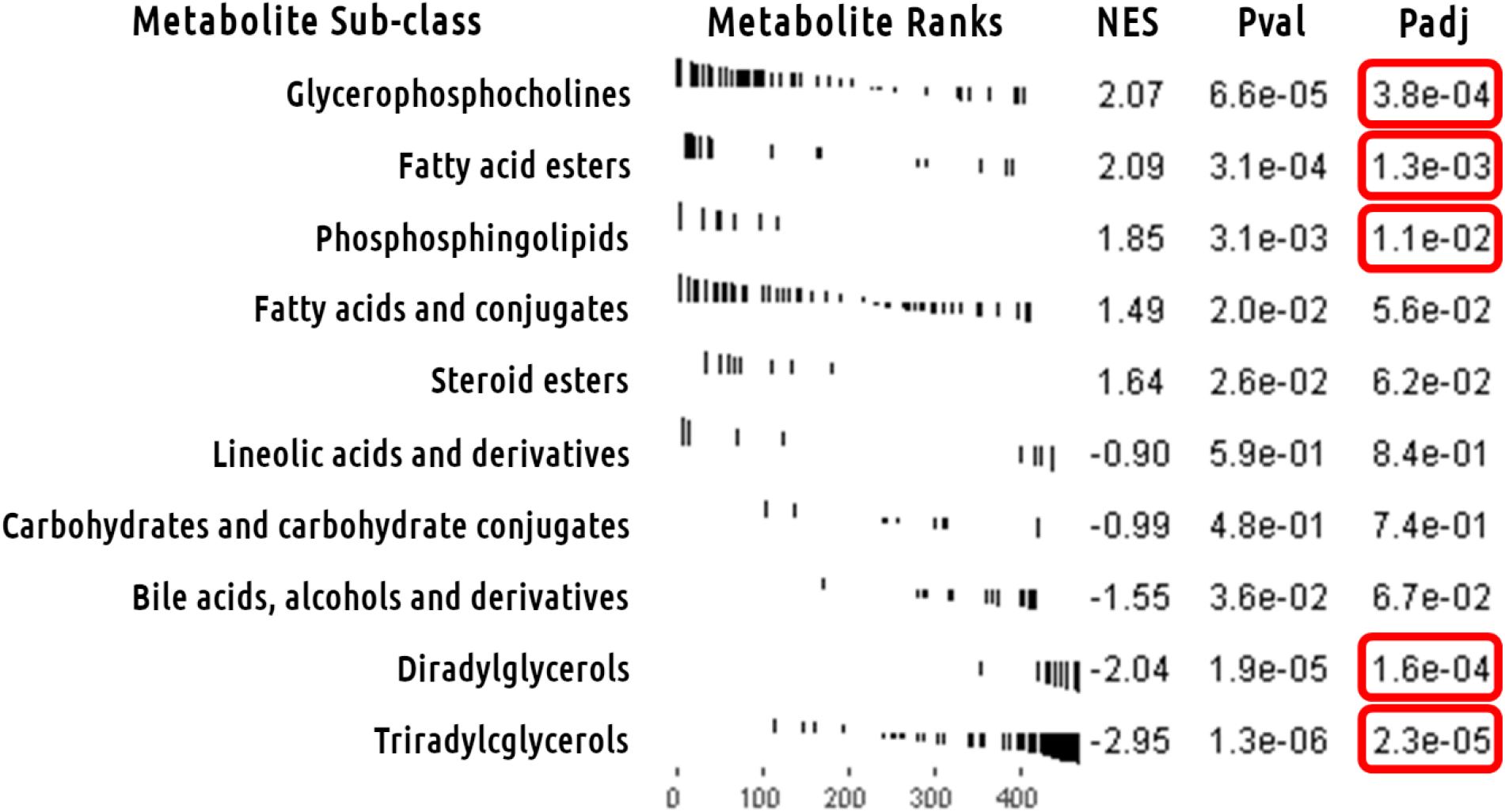
Metabolite Set Analysis of Univariate Model Meta-analysis.

Main metabolite set analysis results for Human Metabolome Database’s (HMDB’s) subclass taxonomic identification of metabolites. Enrichment plots for the top 5 HMDB subclasses associated with increased insulin resistance followed by the top 5 HMDB subclasses associated with reduction in insulin resistance. The x-axis is the metabolome sorted by meta-analysis z-score. The y-axis has a black line for each hit in the *a priori* defined set of metabolites with length equal to the meta-analysis z-score. Triacylglycerols stand out as most significant overall HMDB subclass and most significantly associated with reduction in insulin resistance. “NES” is normalized enrichment score. “Pval” is the p-value for the test. “Padj” is the same p-value after FDR multi-test correction.

### Metabolites showing heterogeneity of effect between different weight loss interventions

In analyses designed to determine which metabolites at baseline associate with heterogeneity in percent change in HOMA-IR between different weight loss intervention types, we performed a Cochran q-test of heterogeneity, with a linear correction for the co-variates of age, sex, race, baseline clinical triglycerides, and change in weight over the intervention time period. This type of analysis is designed to find individual metabolites that can be added to a traditional clinical model to aid in determining which of available obesity intervention a patient should be assigned to. These analyses identified 50 metabolites (Supplemental Table 2) with nominally significant p-values (p<0.05) for heterogeneity by weight loss intervention cohort. Table 3 shows results for the top 10 metabolites from these analyses. The top metabolite, for example, was N6,N6-dimethyllysine (meta-analysis p=1.14e-4), which demonstrated an estimated effect size in percent change of HOMA-IR units of 14.63% in the CBD surgical cohort vs −10.57% in the STRRIDE-PD exercise cohort and −2.04% in the WLM behavioral cohort, suggesting that participants with higher baseline levels of this metabolite have a greater improvement of insulin sensitivity in response to exercise than to surgical weight loss. Figure 4 shows a volcano plot of these results. Unlike the findings in the primary analysis, no major HMDB classification dominates the top results.

**Fig. 4:**
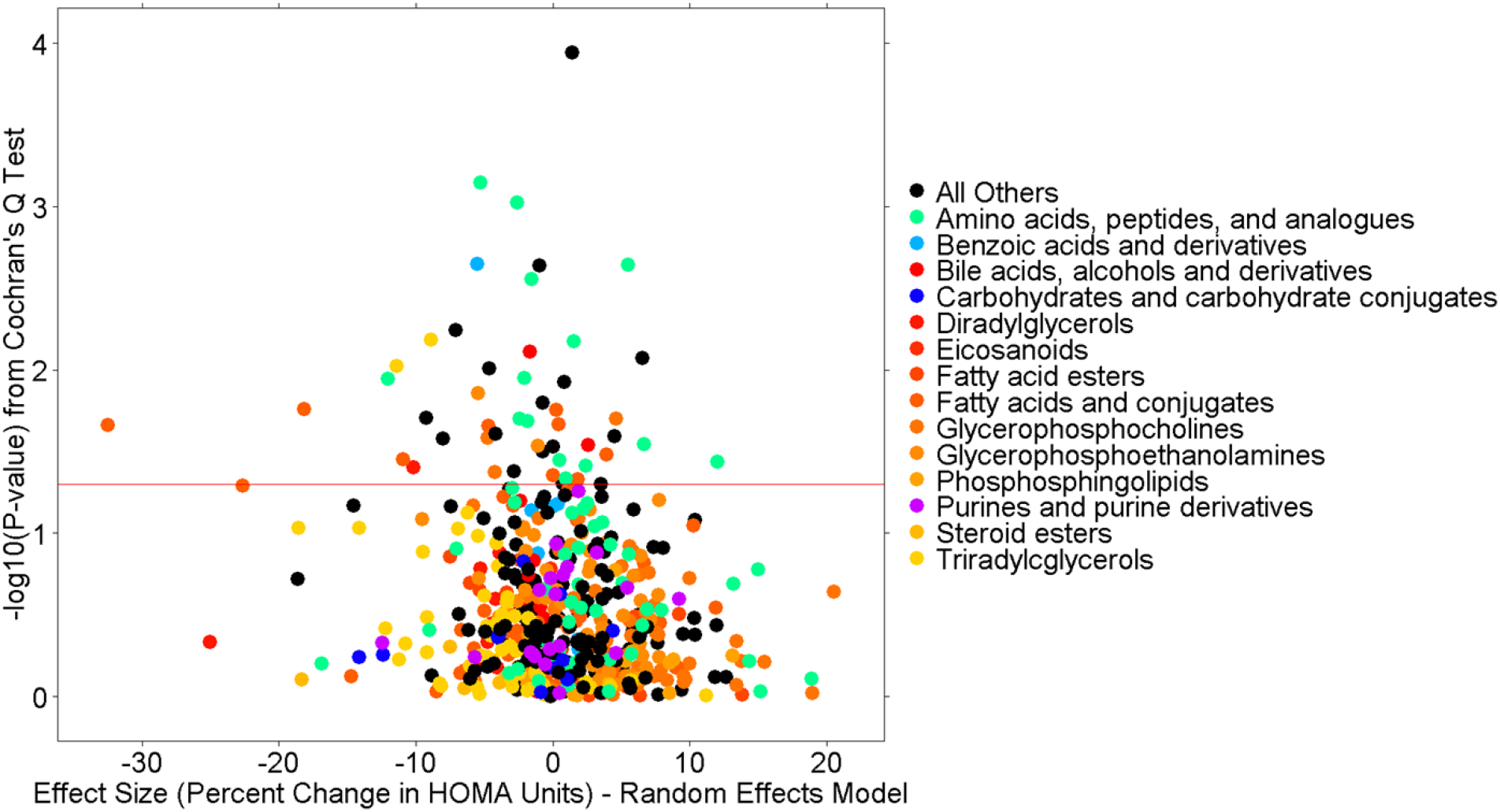
Heterogeneity Volcano Plot from Full Model Meta-analysis.

**Table 3:**
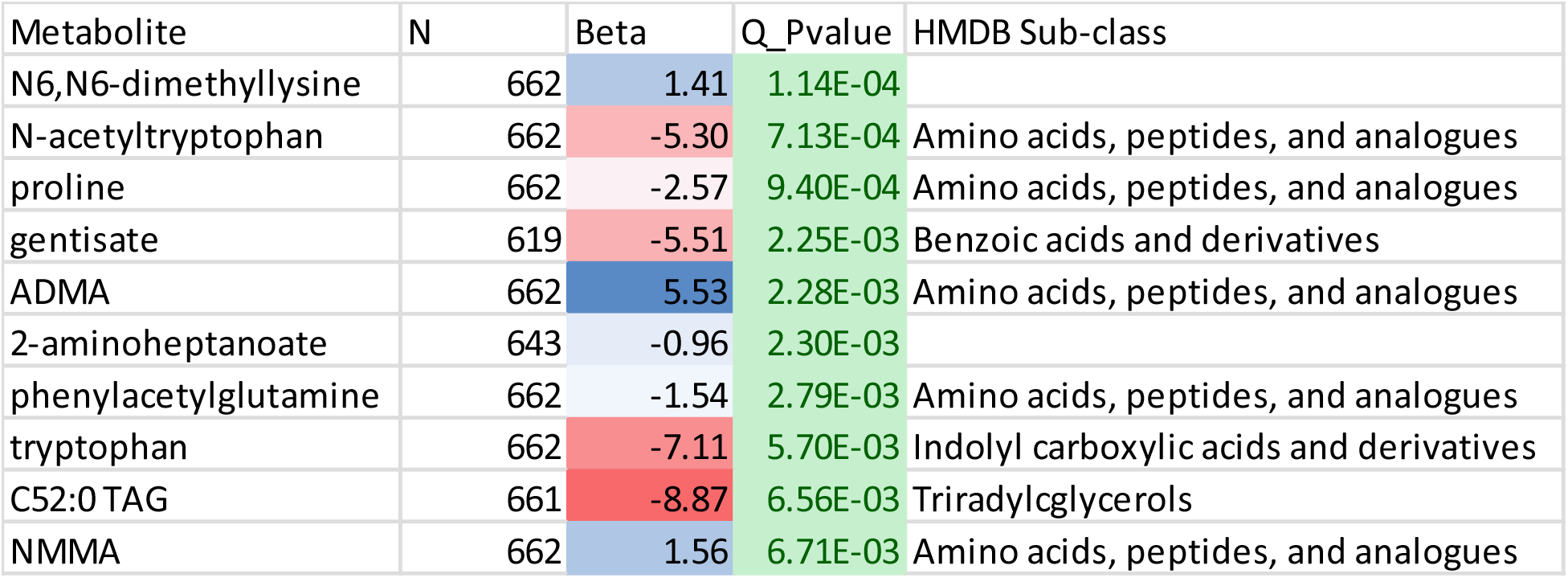
Top 10 Metabolites Heterogeneous in Effect among Interventions from Full Model.

Top 10 metabolites from the heterogeneity analysis of percent change in HOMA-IR. HOMA-IR is Homeostatic Model Assessment of Insulin Resistance. Beta is the random-effects effect-size estimate in percent change of HOMA-IR units (e.g., “−40” indicates a 40% drop in HOMA-IR over the intervention time period). Q_Pvalue is the p-value from the Cochran Q-test of heterogeneity. Subclass is the Human Metabolome Database’s (HMDB’s) subclass taxonomic identification for the metabolite.

Volcano plot of the heterogeneity analysis results from meta-analysis of the three cohorts for the 765 known metabolites from all LC-MS methods in this study. The x-axis is the random-effects effect-size estimate in percent change of HOMA-IR units (e.g., “−40” indicates a 40% drop in HOMA-IR over the intervention time period). Each dot is a metabolite colored by the Human Metabolome Database’s (HMDB’s) subclass taxonomic identification. The y-axis is the – log10(P-value) from the Cochran Q-test of heterogeneity.

### Metabolite set analysis for metabolites associated with heterogeneity of effect among weight loss interventions

Metabolite set analysis was again performed, this time for the set of metabolites associated with heterogeneity in response to intervention using the Cochran q-test statistic. Figure 5 shows the enrichment plots for the top five most heterogeneous followed by the five least heterogeneous sub-classification of metabolites. After FDR multi-test correction for the 22 sub-classifications tested, only the metabolite set “Amino acids, peptides, and analogues” was significant (FDR adjusted p=4.7e-3). This result is not clear from single metabolite analysis and therefore shows the power of metabolite set analyses like this one. Supplemental Table 4 contains results from all tested HMDB sub-classifications.

**Fig. 5:**
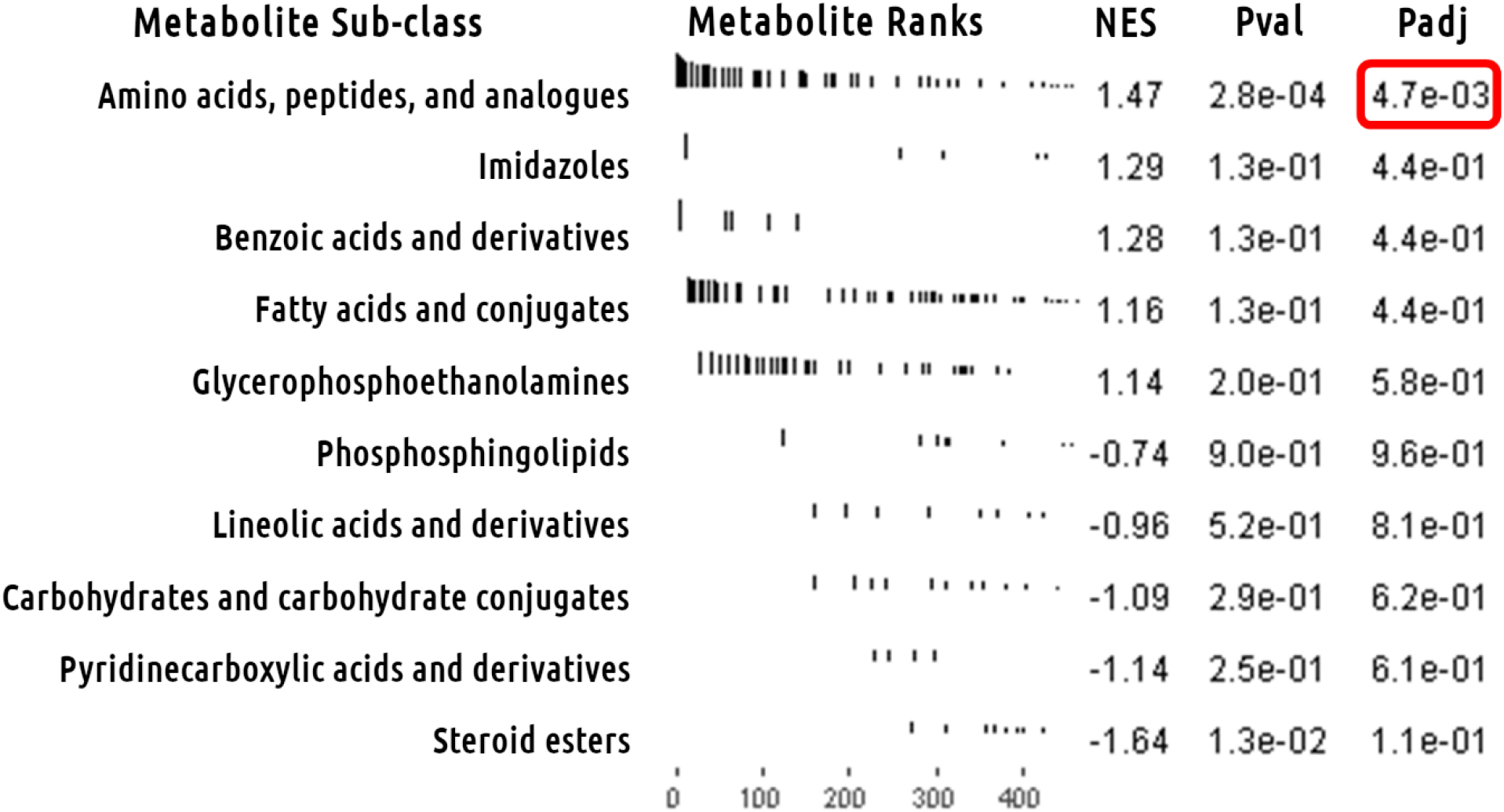
Heterogeneity Metabolite Set Analysis of Full Model Meta-analysis.

Heterogeneity metabolite set analysis results for Human Metabolome Database’s (HMDB’s) subclass taxonomic identification of metabolites. Enrichment plots for top 5 most heterogeneous and top 5 least heterogeneous HMDB subclasses between the three cohorts in this study. The x-axis is the metabolome sorted by heterogeneity Q-score. The y-axis has a black line for each hit in the *a priori* defined set of metabolites with length equal to the heterogeneity Q-score. The amino acids, peptides, and analogues subclass stand out as most significantly heterogeneous HMDB subclass. “NES” is normalized enrichment score. “Pval” is the p-value for the test. “Padj” is the same p-value after FDR multi-test correction.

## Discussion

In the first-of-its-kind study using a comprehensive metabolomics platform in three large cohorts, we have identified metabolites that, measured at baseline, were associated with improvements in insulin resistance, an important metabolic health measure, across behavioral, exercise and surgical weight loss interventions in individuals with obesity. Specifically, we found that higher baseline levels of complex triglyceride lipid species, triacylglycerols and diacylglycerols, are associated with a more salutatory metabolic response across weight loss interventions. Perhaps more importantly, we identify metabolites that were associated with heterogeneity in improvement in insulin resistance depending on the type of weight loss intervention. Specifically, we found 14 amino acids, peptides, and analogues, measured at baseline, that were associated with differential response to weight loss intervention (N-acetyltryptophan, proline, ADMA, phenylacetylglutamine, NMMA, phenylacetylglutamine, tyrosine, hydroxyproline, N-alpha-acetylarginine, N6-acetyllysine, betaine, histidine, 2-aminooctanoate, and lysine; Supplemental Table 2). These metabolites have great potential in precision medicine for overweight/obese individuals, serving as baseline biomarkers that add to clinical models of metabolic response to weight loss interventions, and to help guide a personalized approach to weight loss intervention.

Most dietary fat is TAGs, which need to be broken down before absorption in the gut, then reassembled into circulating low and high-density lipoproteins (LDL/HDL). High levels of TAGs have been associated with atherosclerosis and stroke.[38] DAGs are precursors to TAGs that have themselves be associated with immune-independent mechanisms of developing insulin resistance and/or T2DM in muscle and liver tissues.[39,40]

Previous work has indicated that individual TAGs with lower carbon number (44-52 vs 54-60 carbons) and separately lower double-bond content (0-3 vs 4-12 double-bonds) are associated with a higher likelihood of developing T2DM (higher carbon number and higher double-bond content were neutral in effect towards likelihood of developing T2DM).[41] In the current study, we find individual TAGs with carbon numbers between 50 and 55 and double-bond count between two and three were associated with reduction in insulin resistance over an intervention for obesity time period. Other TAG species were neutral in effect toward insulin resistance or slight reduction in insulin resistance (Supplemental Figure 1 and 2). This may mean low carbon number, saturated TAGs that indicated one may develop T2DM in the previous study[41] may also indicate an obesity intervention will be less effective.

Of note, using a much less comprehensive metabolomic platform, we have previously observed branched chain amino acids (BCAA) to be associated with insulin resistance and that higher baseline levels are associated with a greater decrease in insulin resistance[42,43]. In this study, we find amino acid analogues (including BCAAs) to be heterogeneous among our cohorts. In the case of the individual BCAAs (valine, leucine, isoleucine), they show no association in our CBD surgical cohort, while being positively associated with reduction in insulin resistance in WLM and STRRIDE-PD. The low standard error in the CBD cohort causes the overall inverse-variance weighted meta-analysis of the three cohorts to be non-significant. It is know that gut bacteria can alter the bioavailability of BCAAs.[44] This may indicate that the microbiome is important to consider during exercise/behavioral obesity interventions and less so during surgical interventions perhaps due to antibiotic usage related to having a surgery.

Another item of note is although our top hit of N6,N6-dimethyllysine is relatively unknown in the literature, our second most significantly heterogeneous amino acid analogues, N-acetyltryptophan, demonstrated the same pattern as the BCAAs; it is also been shown to be associated with host-gut microbiota interactions in both blood and urine bio-specimens.[45–47] Namely that the amino acid tryptophan is modified into N-acetyltryptophan by the gut microbiota and then absorbed into the host human. This adds to our earlier statement that the microbiome is important to consider during exercise/behavioral obesity interventions and less so during surgical interventions. We would point to antibiotic usage related to having surgery “resetting” the microbiome after the baseline sample (used in this study to predict outcomes) has been collected as a plausible reason for this.

While this is the first study to compare a large number of metabolites measured at baseline across diverse weight loss interventions, the study has a few limitations. The included cohorts are prospective, but are observational and therefore causation of metabolite pathways on insulin resistance cannot be determined, although we note that these associations remained significant after adjustment for amount of weight loss and other important comorbidities. Further, these results highlight important metabolites that might be used in a prospective randomized biomarker-guided clinical trial of assignment to different weight loss interventions. Our study also primarily involves individuals of European and/or African ancestry and therefore has unclear implications for other ancestries.

We believe this work demonstrates the validity and utility of evaluating the blood metabolome when determining the proper obesity intervention for a patient in a precision medicine context. Our work demonstrates the potential value of measuring individual TAGs and the differentiating ability of amino acid analogues in deciding the best obesity intervention for an individual. Future biomarker-guided intervention studies are necessary to determine clinical utility.

## Acknowledgments

NAB and SHS are funded by American Heart Association Strategically Focused Research Network 17SFRN33670990 and 17SFRN33700155. LCK, CBC, AAD, and SHS are funded by NHLBI 5R01HL127009. REG is funded by NIDDK 5R01DK081572.

## Author contributions

NAB: Data curation, Formal analysis, Investigation, Methodology, Writing

LCK: Data curation

CBC: Methodology

AAD: Methodology

REG: Methodology

NJP: Data curation

BL: Conceptualization, Data curation

LPS: Conceptualization, Data curation

CBN: Data curation, Methodology

WEK: Conceptualization, Data curation

SHS: Conceptualization, Funding acquisition, Project administration, Resources, Supervision, Writing

## Supporting information captions

**Supplemental Table 1: HOMA-IR PC vs logMetabolite - Univariate Model - Analysis** Includes all meta-analysis results for univariate model after filtering (see methods). Metabolite: metabolite name, N_Samples: number of samples, N_Studies: number of cohorts that had data for the metabolite (up to 3), Beta_Fixed: effect size from fixed effects meta-analysis, SE_Fixed: standard error from fixed effects meta-analysis, Zvalue_Fixed: z-score from fixed effects meta-analysis, Pvalue_Fixed: p-value from fixed effects meta-analysis, Beta_Random: effect size from random effects meta-analysis, SE_Random: standard error from random effects meta-analysis, Zvalue_Random: z-score from random effects meta-analysis, Pvalue_Random: p-value from from random effects meta-analysis, Q: Cochran q-test statistic, Q_df: Cochran q-test degrees of freedom, Q_Pvalue: Cochran q-test p-value, Tau2: tau-squared between-study variance, H: H-statistic, I2: I-squared statistic, WLM_GP1_Beta: Effect size within WLM, WLM_GP1_SE: standard error within WLM, WLM_GP1_Num: number of samples within WLM, StrridePD_Beta: effect size within STRRIDE-PD, StrridePD_SE: standard error within STRRIDE-PD, StrridePD_Num: Number of samples within STRRIDE-PD, CBD_Beta: effect size within CBD, CBD_SE: standard error within CBD, CBD_Num: number of samples within CBD, Method: LC-MS method used to measure metabolite, HMDB.ID…representative.ID.: HMDB ID for metabolite, super_class: HMDB metabolite taxonomy super class, class: HMDB metabolite taxonomy class, sub_class: HMDB metabolite taxonomy sub class, missingness: metabolite missingness rate, zeroness: metabolite rate of being zero value, CV: metabolite coefficient of variation, Pvalue_Adj_Fixed: FDR adjusted p-value from fixed effects meta-analysis, Pvalue_Adj_Random: FDR adjusted p-value from random effects meta-analysis.

**Supplemental Table 2: HOMA-IR PC vs logMetabolite - Full Model - Analysis** Includes all meta-analysis results for full model after filtering (see methods). Same columns as Supplemental Table 1.

**Supplemental Table 3: GSEA HOMA-IR PC Taxonomy Subclass Results** Includes all Metabolite Set Analysis of univariate model meta-analysis results for the 22 metabolite sets. Pathway: name of metabolite set, pval: p-value from GSEA test, padj: FDR adjusted p-value, ES: enrichment score, NES: normalized enrichment score, nMoreExtreme: number of 1 million permutation that were more extreme than data, size: number of metabolites in metabolite set.

**Supplemental Table 4: GSEA HOMA-IR PC Heterogeneity Taxonomy Subclass Results** Includes all Heterogeneity Metabolite Set Analysis of full model meta-analysis results for the 22 metabolite sets. Same columns as Supplemental Table 3.

**Supplemental Figure 1: HOMA-IR PC logMetabolite Analysis - Univariate Model - Triradylcglycerols - Carbon Downsloping Plot - Sig** Plot of TAG metabolite effect size from random effects meta-analysis of univariate model vs number of carbon atom in the TAG.

**Supplemental Figure 2: HOMA-IR PC logMetabolite Analysis - Univariate Model - Triradylcglycerols - Bond Downsloping Plot - Sig** Plot of TAG metabolite effect size from random effects meta-analysis of univariate model vs number of double bonds in the TAG.

